# Transcriptomic data reveal divergent paths of chitinase evolution underlying dietary convergence in anteaters and pangolins

**DOI:** 10.1101/2022.11.29.518312

**Authors:** Rémi Allio, Sophie Teullet, Dave Lutgen, Amandine Magdeleine, Rachid Koual, Marie-Ka Tilak, Benoit de Thoisy, Christopher A. Emerling, Tristan Lefébure, Frédéric Delsuc

**Affiliations:** ISEM, Univ. Montpellier, CNRS, IRD, Montpellier, France; CBGP, INRAE, CIRAD, IRD, Montpellier SupAgro, Univ. Montpellier, Montpellier, France; Institute of Ecology and Evolution, University of Bern, Bern, Switzerland; Swiss ornithological Institute, Sempach, Switzerland; Institut Pasteur de la Guyane, Cayenne, French Guiana, France; Kwata NGO, Cayenne, French Guiana, France; Biology Department, Reedley College, Reedley, CA, USA; Université Claude Bernard Lyon 1, LEHNA UMR 5023, CNRS, ENTPE, F-69622, Villeurbanne, France

**Keywords:** Chitinases, Convergent evolution, Myrmecophagy, Mammals, Salivary glands, Transcriptomics

## Abstract

Ant-eating mammals represent a textbook example of convergent evolution. Among them, anteaters and pangolins exhibit the most extreme convergent phenotypes with complete tooth loss, elongated skulls, protruding tongues, hypertrophied salivary glands producing large amounts of saliva, and powerful claws for ripping open ant and termite nests. However, comparative genomic analyses have shown that anteaters and pangolins differ in their chitinase acidic gene (*CHIA*) repertoires, which potentially degrade the chitinous exoskeletons of ingested ants and termites. While the southern tamandua (*Tamandua tetradactyla*) harbors four functional *CHIA* paralogs (*CHIA1*-*4*), Asian pangolins (*Manis* spp.) have only one functional paralog (*CHIA5*). Here, we performed a comparative transcriptomic analysis of salivary glands in 33 placental species, including 16 novel transcriptomes from ant-eating species and close relatives. Our results suggest that salivary glands play an important role in adaptation to an insect-based diet, as expression of different *CHIA* paralogs is observed in insectivorous species. Furthermore, convergently-evolved pangolins and anteaters express different chitinases in their digestive tracts. In the Malayan pangolin, *CHIA5* is overexpressed in all major digestive organs, whereas in the southern tamandua, all four functional paralogs are expressed, at very high levels for *CHIA1* and *CHIA2* in the pancreas, and for *CHIA3* and *CHIA4* in the salivary glands, stomach, liver, and pancreas. Overall, our results demonstrate that divergent molecular mechanisms within the chitinase acidic gene family underlie convergent adaptation to the ant-eating diet in pangolins and anteaters. This study highlights the role of historical contingency and molecular tinkering of the chitin-digestive enzyme toolkit in this classic example of convergent evolution.

## Introduction

Convergent evolution provides a fascinating window into the mechanisms by which similar environmental pressures shape the phenotypes of phylogenetically distant taxa. Indeed, despite the enormous diversity of organisms on Earth and the many potential ways to adapt to similar conditions, the strong deterministic force of natural selection has led to numerous instances of recurrent phenotypic adaptation (Losos 2011; McGhee 2011; Losos 2018).

Although classical models of convergence at the molecular level often assume identical mutations in the same genes across species (Arendt and Reznick 2008), emerging evidence from comparative genomics and transcriptomics suggests that the recruitment of the same or similar genes and pathways may also lead to similar phenotypes across divergent lineages. For instance, convergent electric fish, which have evolved independently at least six times, provide a good illustration of the complexity of the selective process that follows from the interaction of contingency, constraints, and convergence (Zakon et al. 2006). In this case, the same genes have been independently recruited and differentially expressed in novel electric organs due to developmental constraints, and their function subsequently adjusted by natural selection involving convergent amino acid substitutions in functionally important domains (Galant et al. 2014; Liu et al. 2019; Wang & Yang 2021). This suggests an important role for evolutionary constraints imposed by existing genomic architectures and developmental pathways, leading to the repeated use of similar genetic material in the origin of evolutionary novelties (Shubin et al. 2009). In this context, historical contingency often leads to evolutionary tinkering as natural selection works from available material (Jacob 1977). Thus, both historical contingency and deterministic evolution appear to have influenced the evolution of current biodiversity, and one of the key questions is to assess the relative influence of these two evolutionary processes (Blount et al. 2018).

As intuited by Jacob (1977), molecular tinkering appears to be particularly common and has indeed shaped the evolutionary history of a number of gene families (McGlothlin et al. 2016; Pillai et al. 2020; Xie et al. 2021). The particular evolutionary dynamics observed in gene families can lead to both evolutionary opportunities due to gene duplications paired with the acquisition of a new function but also evolutionary constraints due to ancestral loss of function. A good example resides in the evolution of chitinase genes in placental mammals, which belong to the large Glycosyl Hydrolase 18 (GH18) gene family (Bussink et al. 2007; Funkhouser and Aronson 2007). Recent studies have shown that chitinase genes may play an important digestive function in insectivorous species (Emerling et al. 2018; Janiak et al. 2018; Wang et al. 2020; Cheng et al. 2022). Indeed, while the placental ancestor possessed five functional chitinase acidic (*CHIA*) paralogs, the evolution of this gene family was subsequently shaped through multiple pseudogenization events associated with dietary adaptation during the placental radiation (Emerling et al. 2018). Interestingly, the widespread gene loss observed in carnivorous and herbivorous lineages in particular, resulted in a global positive correlation between the number of functional *CHIA* paralogs and the percentage of invertebrates in the diet across placentals. Indeed, mammals with a low proportion of insects in their diet present none or only a few functional *CHIA* paralogs and those with a high proportion of insects in their diet generally have retained four or five functional *CHIA* paralogs (Emerling et al. 2018; Janiak et al. 2018; Wang et al. 2020; Fig. 1).

**Figure 1:**
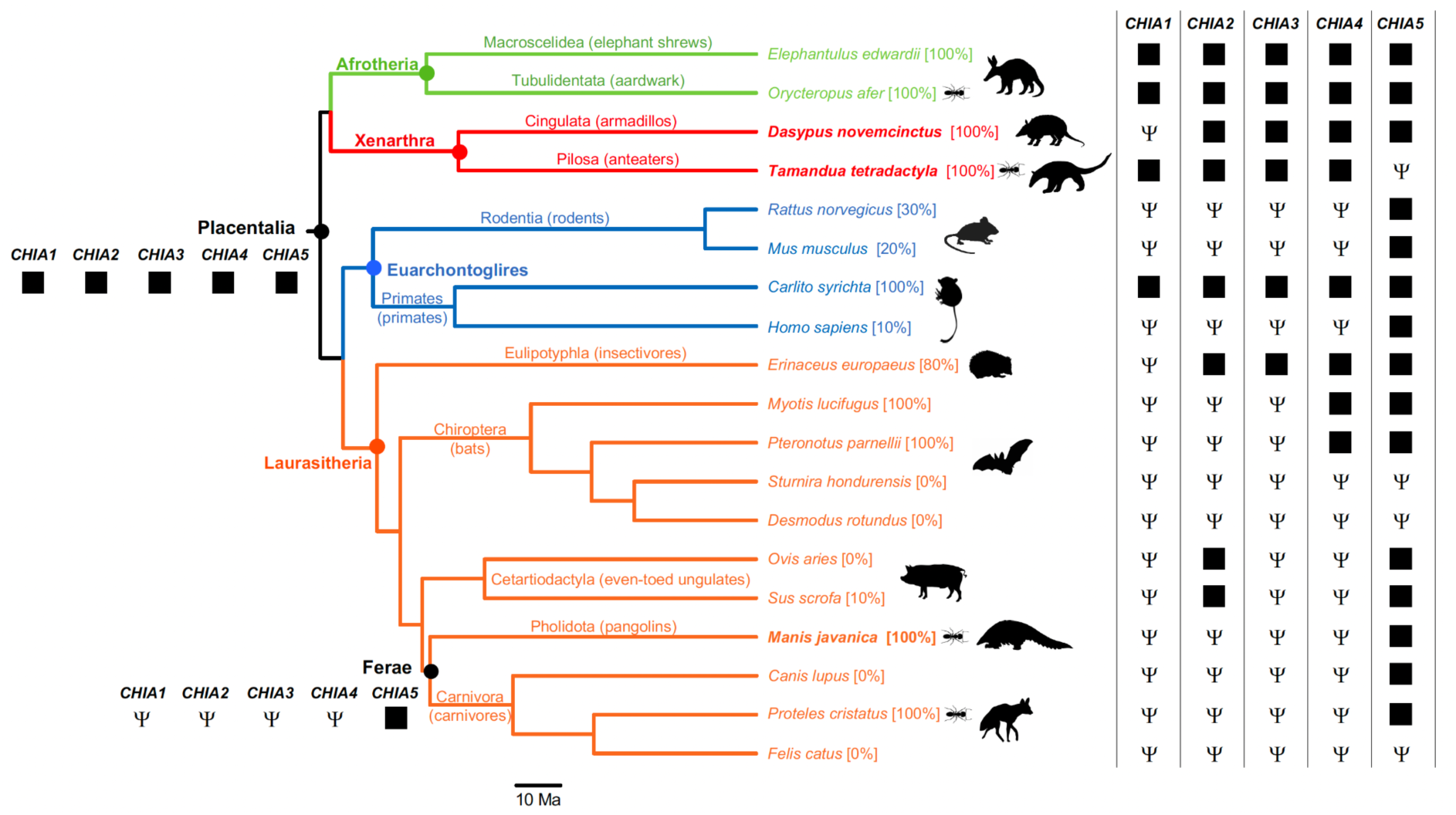
Dated placental mammal phylogeny including representative species of the four major clades (Afrotheria, Xenarthra, Euarchontoglires, and Laurasiatheria) for which *CHIA* gene repertoires have been previously characterized. Numbers between brackets represent percentages of invertebrates included in the diet with myrmecophagous species indicated by an ant silhouette. Ψ symbols indicate *CHIA* pseudogenes as determined in previous studies (Emerling et al. 2018; Janiak et al. 2018; Wang et al. 2020). Ancestral *CHIA* gene repertoires for Placentalia and Ferae (Pholidota + Carnivora) as inferred by Emerling et al. (2018) are presented. The chronogram was extracted from www.timetree.org (Kumar et al. 2022). Silhouettes were obtained from www.phylopic.org.

Myrmecophagous mammals, with more than 90% of their diet consisting of social insects (Redford 1987), have convergently evolved dietary adaptation such as powerful claws used to dig into ant and termite nests, tooth reduction culminating in complete tooth loss in anteaters and pangolins (Ferreira-Cardoso et al. 2019), an elongated muzzle with an extensible tongue (Ferreira-Cardoso et al. 2020), and viscous saliva produced by hypertrophied salivary glands (Reiss 2001). With regards to their chitinase gene repertoire, they are generally grouped with the most insectivorous species (Fig. 1). Specifically, the southern tamandua (*Tamandua tetradactyla*) and the aardvark (*Orycteropus afer*) indeed possess four (*CHIA1-4*) and five (*CHIA1-5*) functional paralogs, respectively. However, pangolins appear as a striking exception. Despite their strict myrmecophagous diet and many associated convergent features shared with other myrmecophagous species (anteaters in particular), the two investigated species (*Manis javanica* and *M. pentadactyla*) possess only one functional *CHIA* paralog (*CHIA5*). The presence of the sole *CHIA5* in pangolins was hypothesized to be the consequence of historical contingency on the evolution of the chitinase family with the probable loss of *CHIA1-4* functionality in the almost recent common ancestor of Pholidota and Carnivora (Ferae; Emerling et al. 2018; Fig. 1). It has indeed recently been confirmed that a non insect-based diet has caused structural and functional changes in the *CHIA* gene repertoire resulting in multiple losses of function in Carnivora with only few species including insects in their diet retaining a fully functional *CHIA5* gene (Tabata et al. 2022). These recent results, combined with the apparent importance of chitinase paralogs in insect digestion, have prompted questions regarding how pangolins succeed in digesting chitin with only one functional paralog.

One possible evolutionary solution for inheriting a depleted gene family resides in the modification of gene expression patterns in the remaining functional paralogs. Indeed, *CHIA5* was recently found to be highly expressed in the main digestive organs of the Malayan pangolin (Ma et al. 2017; Ma et al. 2019; Cheng et al. 2023) suggesting that pangolins might compensate for their reduced chitinase repertoire by an increased ubiquitous expression of their only remaining functional *CHIA5* paralog in multiple organs. While this result is very encouraging, it lacks a general comparison with *CHIA* paralogs expression in other mammals and more specifically with other myrmecophagous mammals that present more functional *CHIA* paralogs. If gene expression indeed plays a compensatory role, one can expect that *CHIA5* expression in pangolins would be comparatively higher and more ubiquitous among digestive organs than the expression of the other *CHIA* paralogs in convergent myrmecophagous species.

To further explore *CHIA* paralog expression in mammals and more particularly in convergent myrmecophagous species, we adopted a threefold approach. First, with the aim of identifying all functional paralogs and better understanding their function in chitin digestion, we reconstructed the first detailed evolutionary history of the chitinase-like gene family in mammals based on phylogenetic analysis of publicly available genomic and transcriptomic data. In a second step, we generated a large comparative dataset of salivary gland transcriptomes encompassing 33 mammalian species from various lineages with diverse diets (herbivores, carnivores, frugivores, insectivores, omnivores), enabling for the first time the comparison of *CHIA* expression across mammalian species. The objective here was to determine whether insectivores and myrmecophagous species indeed exhibit differential chitinase paralog expression in their salivary glands compared to mammals with other diets. In a third step, we focused on two convergent myrmecophagous species (the southern tamandua and the Malayan pangolin) and an insectivorous species (the nine-banded armadillo) for which we were able to assemble and generate transcriptomes of several digestive and non-digestive tissues, to compare the use of their chitinase gene repertoire expression across different organs. The objective of this final step was to determine whether variations in genomic chitinase repertoires were associated with distinct expression patterns in digestive tissues or whether these patterns were independent of the functional gene repertoire. Overall, by leveraging species diversity on the one hand and organ diversity on the other, our results shed light on the molecular underpinnings of convergent evolution in ant-eating mammals by revealing that divergent paths of chitinase gene family evolution underlie dietary convergence between anteaters and pangolins.

## Results

### Mammalian chitinase gene family evolution

In order to gain further insights into the evolution and potential function of chitinase-related genes in mammalian genomes, we performed the first detailed phylogenetic reconstruction of the chitinase-like gene family based on functional paralogs using a gene tree/species tree reconciliation approach. The reconciled maximum likelihood tree of mammalian chitinase genes is presented in Fig. 2A. Our analyses showed that this gene family is constituted by nine paralogs whose evolution is notably characterized by gene loss with 384 speciation events followed by gene loss and 48 gene duplications as estimated by the gene tree/species tree reconciliation algorithm of GeneRax. At the base of the reconciled gene tree, we found the clade *CHIA1-2/OVGP1* (optimal root inferred by the reconciliation performed with TreeRecs) followed by a duplication separating the *CHIT1/CHI3L1-2* and *CHIA3-5* groups of paralogs. Within the *CHIT1/CHI3L* clade, two consecutive duplications gave rise to *CHIT1*, then *CHI3L1* and *CHI3L2*. In the *CHIA3-5* clade, a first duplication separated *CHIA3* from *CHIA4* and *CHIA5*, which were duplicated subsequently. Marsupial *CHIA4* sequences were located at the base of the *CHIA4-5* clade suggesting that this duplication might be recent and specific to placentals. This scenario of chitinase gene evolution is consistent with our new synteny analysis showing physical proximity of *CHIA1-2* and *OVGP1* on one hand, and *CHIA3-5* on the other hand (Fig. 2B), which implies that chitinase genes evolved by successive tandem duplications. However, evidence of gene conversion between the two more recent duplicates (*CHIA4* and *CHIA5*) at least in some taxa suggests that further data are necessary to fully disentangle the origins of these two paralogs.

**Figure 2:**
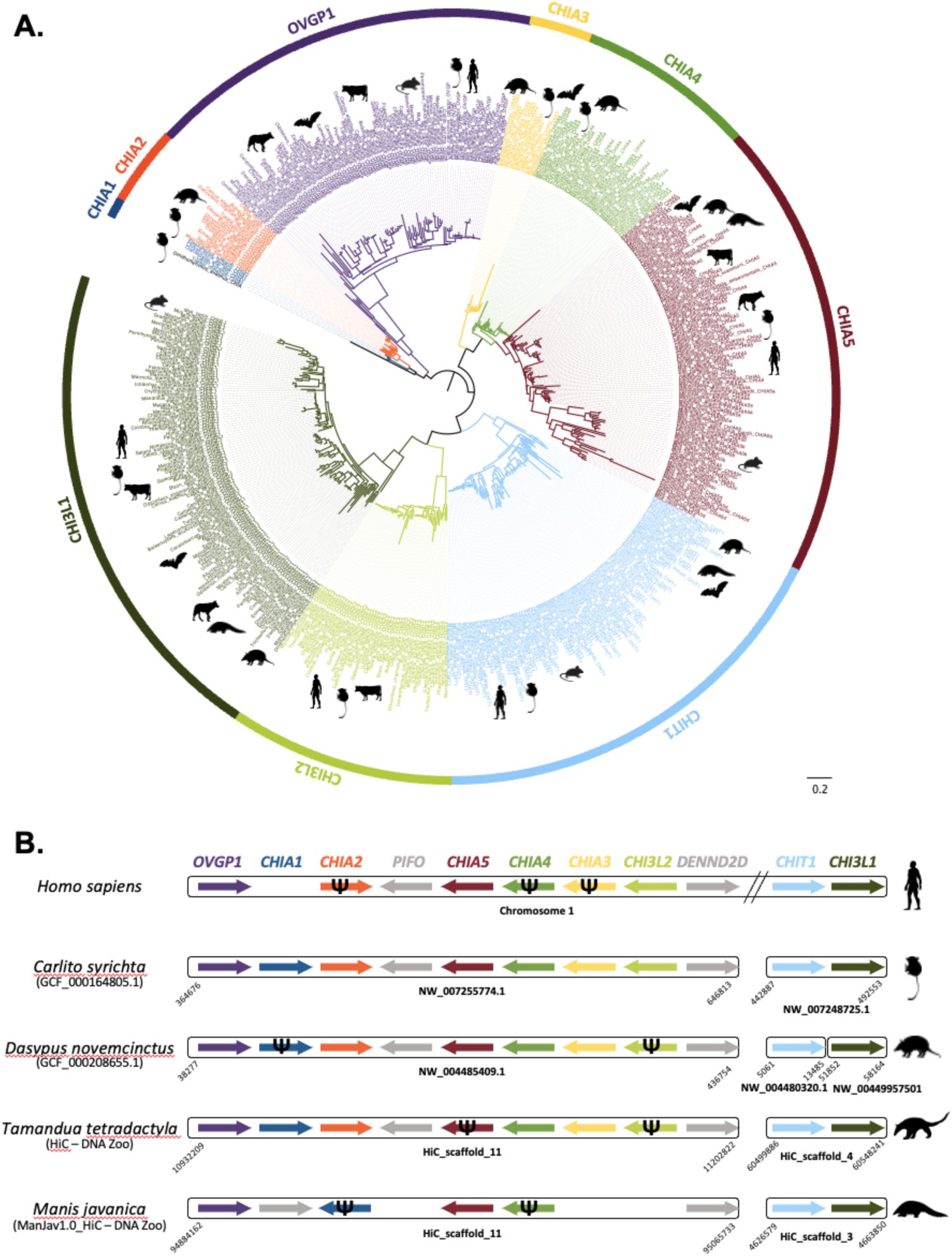
A. Mammalian chitinase-like gene family tree reconstructed using a maximum likelihood gene-tree/species-tree reconciliation approach on protein sequences. The nine chitinase paralogs are indicated on the outer circle. Scale bar represents the mean number of amino acid substitutions per site. B. Synteny analysis of the nine chitinase paralogs in humans (*Homo sapiens*), tarsier (*Carlito syrichta*), nine-banded armadillo (*Dasypus novemcinctus*) and the two main focal convergent ant-eating species: the southern tamandua (*Tamandua tetradactyla*) and the Malayan pangolin (*Manis javanica*). Assembly names and accession numbers are indicated below species names. Boxes represent different contigs with their most upstream and downstream BLAST hit positions to chitinase genes (colored arrows). Genes *PIFO* and *DENND2D* (grey arrows) are not chitinase paralogs but were used in the synteny analysis. Arrow direction indicates gene transcription direction as inferred in Genomicus v100.01 (Nguyen et al. 2022) for genes located on short contigs. Ψ symbols indicate pseudogenes as determined in Emerling et al. (2018). Genes with non significant BLAST hits were not represented and are probably not functional or absent. Silhouettes were obtained from www.phylopic.org.

### Comparison of ancestral sequences

The ancestral amino acid sequences of the nine chitinase paralogs were reconstructed from the reconciled mammalian gene tree and compared to gain further insight into the potential function of the enzymes they encode (Fig. 3; Complete ancestral sequences and associated probabilities available from Zenodo). The alignment of predicted amino acid sequences locates the chitinolytic domain between positions 133 and 140 with the preserved pattern DXXDXDXE. The ancestral sequences of CHI3L1 and CHI3L2, as all contemporary protein sequences of the corresponding genes, have a mutated chitinolytic domain with absence of a glutamic acid at position 140 (Fig. 3A), which is the active proton-donor site necessary for chitin hydrolysis (Olland et al. 2009; Hamid et al. 2013). This indicates that the ability to degrade chitin has likely been lost before the duplication leading to CHI3L1 and CHI3L2 (Fig. 3B). The ancestral sequence of OVGP1 also presents a mutated chitinolytic site although the glutamic acid in position 140 is present (Fig. 3A). The evolution of the different chitinases therefore seems to be related to changes in their active site. The six cysteine residues allowing the binding to chitin are found at positions 371, 418, 445, 455, 457 and 458 (Fig. 3C). The absence of one of these cysteines prevents binding to chitin (Tjoelker et al., 2000) as this is the case in the ancestral OVGP1 protein where the last four cysteine residues are changed (Fig. 3C). The other ancestral sequences present the six conserved cysteine residues and thus can bind to chitin (Fig. 3C).

**Figure 3:**
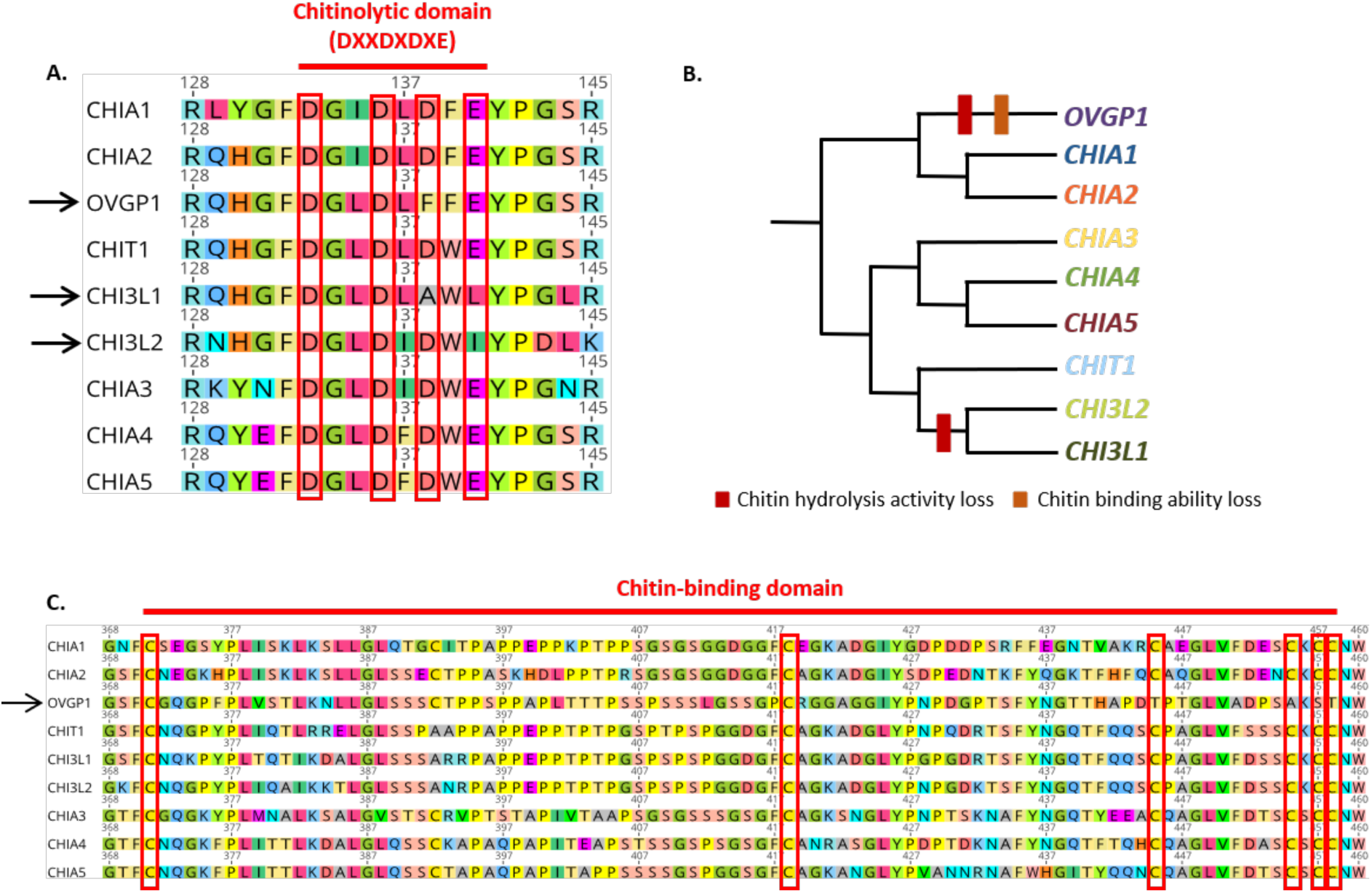
Comparison of predicted ancestral protein sequences of the nine mammalian chitinase paralogs. A. Conserved amino acid residues of the canonical chitinolytic domain active site (DXXDXDXE). Arrows indicate paralogs in which changes occurred in the active site. B. Summary of the evolution of chitinase paralogs functionality. C. Conserved cysteine residues of the chitin-binding domain. The arrow indicates OVGP1 in which the last four cysteines have been replaced.

### Chitinase gene expression in mammalian salivary glands

To test the hypothesis that salivary glands play an important functional role in the digestion of ants and termites in ant-eating mammals, we analyzed the gene expression profiles of the nine chitinase paralogs revealed by the gene family tree reconstruction in 40 salivary gland transcriptomes representing 33 species (Fig. 4). *CHIA1* was expressed only in the elephant shrew (*Elephantulus myurus*; 23.22 normalized read counts [NC]). *CHIA2* was expressed only in the wild boar (*Sus scrofa*; 48.84 NC). *CHIA3* was expressed in the two insectivorous California leaf-nosed bat individuals (*Macrotus californicus*; 367.70, and 35.03 NC) and in all three southern tamandua individuals (*T. tetradactyla*; 48.66, 41.52, and 15.14 NC). *CHIA4* was also highly expressed in all three southern tamanduas (565.61, 214.83, and 180.26 NC) and in the two California leaf-nosed bats (*M. californicus*; 17,224.06, and 16,880.24 NC), but also in the giant anteater (*M. tridactyla*; 50.74 NC). Expression of *CHIA5* was at least an order of magnitude higher in the two Malayan pangolin individuals (*Manis javanica*; 196,778.69 and 729.18 NC) and Thomas’s nectar bat (*Hsunycteris thomasi*; 7,301.82 NC) than in the three other species in which we detected expression of this gene: the domestic mouse (*Mus musculus*; 40.15 NC), common genet (*Genetta genetta*; 132.64 NC), and wild boar (*S. scrofa*; 152.20 NC). *CHIT1* was expressed in many species (12 out of 40 samples) with values ranging from 46.76 NC in a single southern tamandua (*T. tetradactyla*) individual to 115,739.25 NC in the short-tailed shrew tenrec (*Microgale brevicaudata*). *CHI3L1* was expressed in most species (24 out of 40 samples) with values ranging from 61.68 NC in the giant anteater (*M. tridactyla*) to 1,297.01 NC in a Malayan pangolin (*M. javanica*) individual. *CHI3L2* was expressed in human (*H. sapiens*; 1334.07 NC), wild boar (*S. scrofa*; 246.41 NC), elephant shrew (*E. myurus*; 94.65 NC), and common tenrec (*Tenrec ecaudatus*; 68.62 NC). *OVGP1* was only found expressed at very low levels in domestic dog (*Canis lupus familiaris*; 6.80 NC), human (*H. sapiens*; 15.33 NC), one of the two Malayan pangolins (*M. javanica*; 4.99 NC) and wild boar (*S. scrofa*; 17.84 NC). Finally, the southern aardwolf (*P. cristatus*), Norway rat (*Rattus norvegicus*), Parnell’s mustached bat (*Pteronotus parnellii*) and six phyllostomid bat species (*Carollia sowelli*, *Centurio senex*, *Glossophaga commissarisi*, *Sturnira hondurensis*, *Trachops cirrhosus*, and *Uroderma bilobatum*) did not appear to express any of the nine chitinase gene paralogs in any of our salivary gland samples.

**Figure 4:**
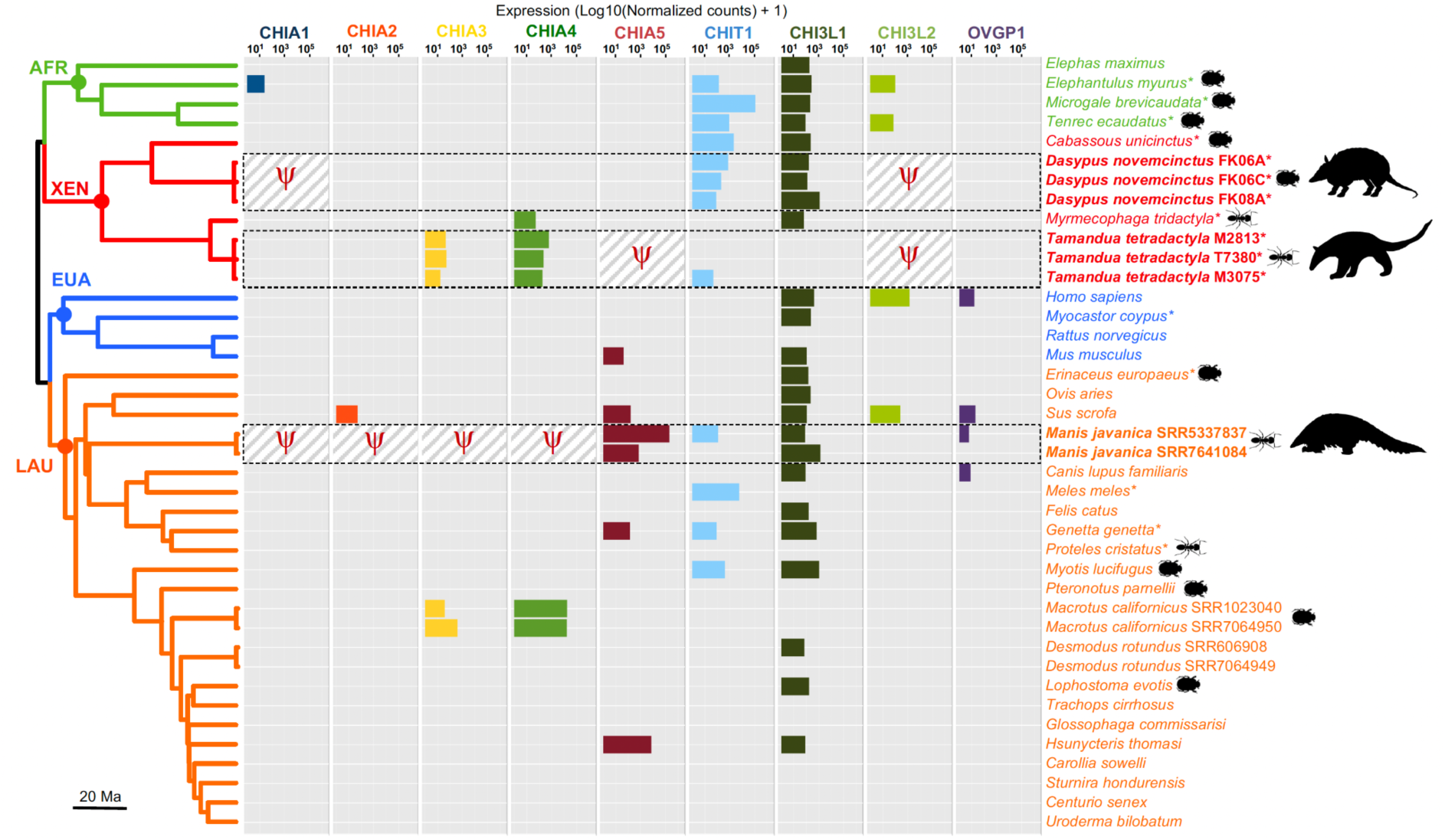
Expression of the nine chitinase paralogs in 40 mammalian salivary gland transcriptomes. The 33 species are presented in their phylogenetic context covering the four major placental clades: Afrotheria (AFR), Xenarthra (XEN), Euarchontoglires (EUA), and Laurasiatheria (LAU). The chronogram was extracted from www.timetree.org (Kumar et al. 2022). Non-functional pseudogenes are only indicated for the three focal species (in bold) using a Ψ symbol: nine-banded armadillo (*Dasypus novemcinctus*), southern tamandua (*Tamandua tetradactyla*) and Malayan pangolin (*Manis javanica*). Expression level is represented as log10 (Normalized Counts + 1). Asterisks indicate the 16 new transcriptomes produced in this study. Myrmecophagous and insectivorous species are indicated by ant and beetle silhouettes, respectively. Silhouettes were obtained from www.phylopic.org.

### Chitinase gene expression in digestive and non-digestive organs

The expression level of the nine chitinase paralogs in several organs was compared among three species including an insectivorous xenarthran (the nine-banded armadillo; *D. novemcinctus*) and two of the main convergent myrmecophagous species (the southern anteater; *T. tetradactyla*, and the Malayan pangolin; *M. javanica*). This analysis revealed marked differences in expression level of these genes among the three species and among their digestive and non-digestive organs (Fig. 5). In the nine-banded armadillo (*D. novemcinctus*), although only *CHIA1* is pseudogenized and consequently not expressed, we did not detect any expression of *CHIA2*, *CHIA3*, and *CHIA4* in the tissues studied here, and *CHIA5* was only weakly expressed in one spleen sample (51.90 NC). In the Malayan pangolin (*M. javanica*), whereas *CHIA1-4* are non-functional and consequently not expressed, *CHIA5* was found expressed in all digestive organs with particularly high levels in the stomach (377,324.73 and 735,264.20 NC) and salivary glands (196,778.69 and 729.18 NC), and at milder levels in the tongue (121.24 NC), liver (254.79 NC on average when expressed), pancreas (168.64 and 39.33 NC), large intestine (238.45 and 79.32 NC), and small intestine (847.51 and 13.72 NC), but also in skin (178.95 NC) and spleen (12.06 NC) samples. Conversely, in the southern tamandua (*T. tetradactyla*), only *CHIA5* is pseudogenized and accordingly not expressed (Fig. 5). *CHIA1* was found highly expressed in the pancreas (64,443.05 NC) and weakly expressed in testes (22.74 and 14.73 NC), and *CHIA2* also had very high expression in the pancreas (1,589,834.39 NC), and low expression in testes (36.51 and 34.52 NC) and lungs (8.22 NC). *CHIA3* was also expressed in the pancreas (359.03 NC), testes (241.79 and 35.42 NC), tongue (39.53 and 12.44 NC), salivary glands (48.66, 41.52, and 15.14 NC), and liver (32.40 NC). Finally, *CHIA4* was expressed in the testes (19.48 and 14.59 NC), spleen (109.97 and 73.31 NC), lungs (340.84 NC), salivary glands (565.61, 214.83, and 180.26 NC), and glandular stomach (116.11 NC). More globally, *CHIT1* was expressed in all tissues in *M. javanica,* in the testes, tongue, salivary glands, and small intestine in *T. tetradactyla*, and in the cerebellum, lungs, salivary glands, and liver in *D. novemcinctus*. *CHI3L1* was found to be expressed in the majority of digestive and non-digestive tissues in all three species. *CHI3L2* is non-functional or even absent in the genome of these three species and was consequently not expressed. *OVGP1* was only weakly expressed in the lungs and salivary glands of *M. javanica* (2.22 and 4.99 NC, respectively).

**Figure 5:**
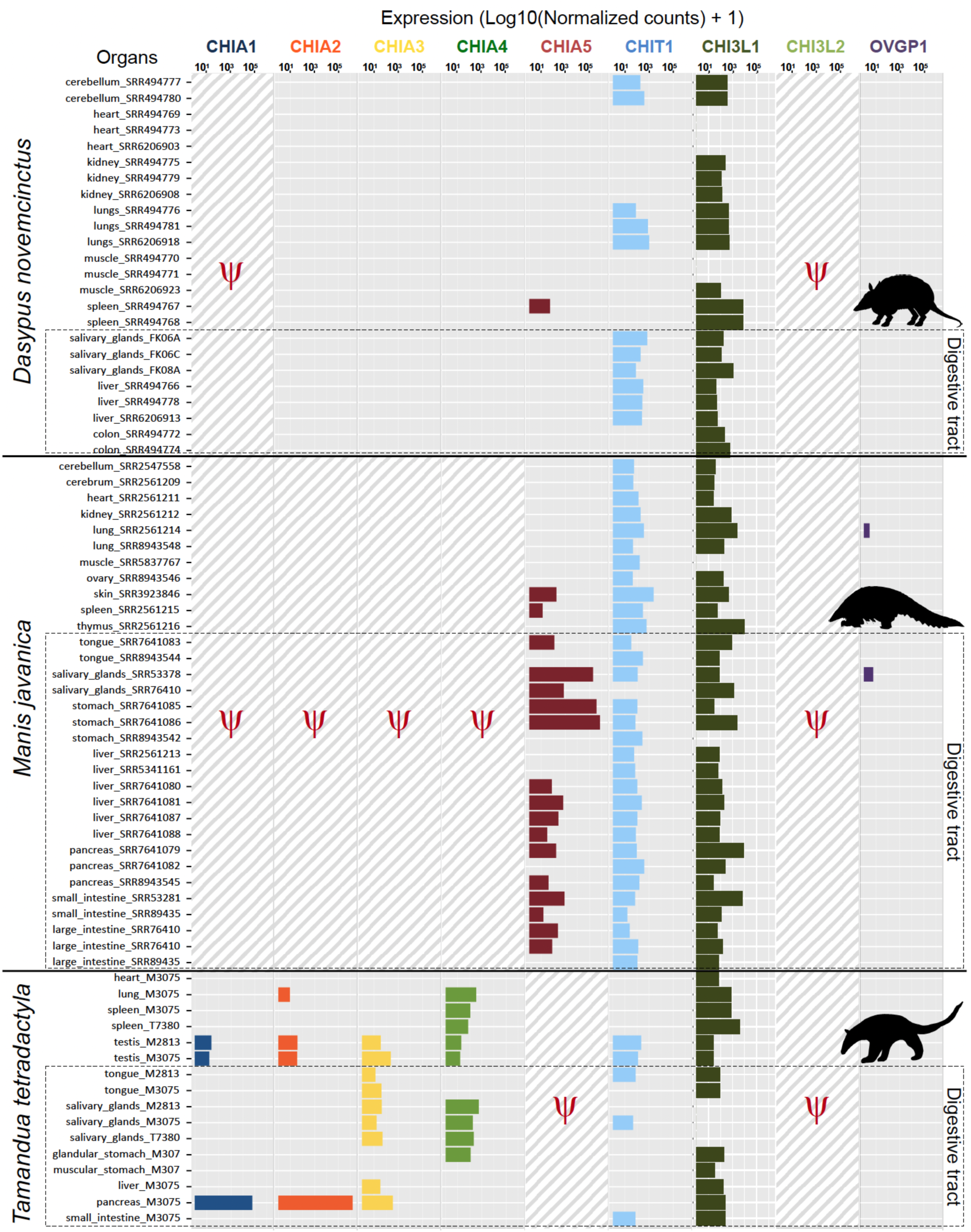
Expression of the nine chitinase paralogs in 72 transcriptomes from different organs of the three focal species: the nine-banded armadillo (*Dasypus novemcinctus*), the Malayan pangolin (*Manis javanica*), and the southern tamandua (*Tamandua tetradactyla*). Non-functional pseudogenes are represented by a Ψ symbol and hatched background. Boxes indicate organs of the digestive tract. Expression level is represented as log10 (Normalized Counts + 1). Silhouettes were obtained from www.phylopic.org.

## Discussion

### Evolution of chitinase paralogs towards different functions

Chitinases have long been suggested to play an important role in insect digestion within mammals (Jeuniaux 1961; Jeuniaux 1966; Jeuniaux 1971; Jeuniaux and Cornelius 1997). After the initial discovery of a single chitinase gene (Boot et al. 2001), comparative genomics and phylogenetics have revealed a gene family (Glycosyl Hydrolase Family 18, GH18) in which chitinases and chitinase-like proteins may work together to facilitate chitin digestion in the digestive tracts of mammals. The first phylogenetic analyses of this gene family have revealed a dynamic evolutionary history and a high degree of synteny among mammals (Bussink et al. 2007; Hussain and Wilson 2013). Our new exhaustive maximum likelihood phylogenetic analyses recovered nine functional paralogous chitinase gene sequences in mammalian genomes (Fig. 2A). In addition to the five previously characterized *CHIA* paralogs (Emerling et al. 2018; Janiak et al. 2018), we were able to identify an additional gene, *OVGP1*, which is most closely related to the previously characterized *CHIA1* and *CHIA2* genes. In placentals, OVGP1 plays a role in fertilization and embryonic development (Buhi 2002; Saint-Dizier et al. 2014; Algarra et al. 2016; Laheri et al. 2018). However, other aliases for OVGP1 include Mucin 9 and CHIT5 suggesting a possible digestive function. This result was further confirmed by synteny analyses suggesting a common origin by tandem duplication for *CHIA1-2* and *OVGP1* within the conserved chromosomal cluster that also includes *CHIA3-5* and *CHI3L2* (Fig. 2B). Comparison of the ancestral amino acid sequences of the nine chitinase paralogs revealed differences in their ability to bind and degrade chitin (Fig. 3), suggesting that these paralogs have evolved towards different functional specializations. The evolution of chitinase-like proteins was accompanied by a loss of enzymatic activity for chitin hydrolysis, which occurred several times independently (Bussink et al. 2007; Funkhouser and Aronson 2007; Hussain and Wilson 2013; Fig. 3B).

CHI3L1 and CHI3L2, which are expressed in various cell types including macrophages and synovial cells, play roles in cell proliferation and immune response (Recklies et al. 2002; Areshkov et al. 2011; Lee et al. 2011). In contrast to these chitinase-like proteins, CHIT1 and the five CHIAs are able to degrade chitin. In humans, *CHIT1* is expressed in macrophages and neutrophils and is suspected to be involved in the defense against chitin-containing pathogens such as fungi (Gordon-Thomson et al. 2009; Lee et al. 2011). In addition to their role in chitin digestion (Boot et al. 2001), CHIAs are also suggested to play a role in the inflammatory response (Lee et al. 2011) and are expressed in non-digestive tissues, in agreement with our comparative transcriptomic results. Thus, it has been proposed that the expansion of the chitinase gene family is related to the emergence of the innate and adaptive immune systems in vertebrates (Funkhouser and Aronson 2007).

The evolution of the different *CHIA1-5* genes has involved changes in their catalytic sites, which have consequences for the secondary structure of enzymes and potentially affect their optimal pH or function, as it has recently been shown for *CHIA5* in Carnivora (Tabata et al. 2022). Experimental testing of the chitin degrading activity of ancestral reconstructions of each of the five CHIA enzymes, on different substrates and at different pH of enzymes, would help determine if there are differences in organ specificity of each enzyme.

Furthermore, studying the potential molecular binding properties of these enzymes to other substrates would shed additional light on their functional roles. For example, changing a cysteine in the chitin-binding domain prevents binding to this substrate but not to tri-N-acetyl-chitotriose (Tjoelker et al. 2000), a compound derived from chitin with antioxidant properties (Chen et al. 2003; Salgaonkar et al. 2015). Such functional assays, complemented by transcriptomic data to determine their expression profile in different tissues and organs (as previously done in the Malayan pangolin; Yusoff et al. 2016; Ma et al. 2017; Ma et al. 2019; Cheng et al. 2023), may help to decipher their respective roles in mammalian digestion (see below).

### Impact of historical contingency and molecular tinkering on chitinase evolution and expression

In the specific case of adaptation to myrmecophagy, comparative genomic and transcriptomic analyses of these chitinase genes, particularly those encoding chitinolytic enzymes (*CHIA*s), have led to a better understanding of how convergent adaptation to myrmecophagy in placentals occurs at the molecular level (Emerling et al. 2018; Cheng et al. 2022). On the one hand, anteaters (Pilosa; Vermilingua) likely inherited five *CHIA* genes from an insectivorous ancestor (Emerling et al. 2018), but then the *CHIA5* gene was lost at least in some of its descendants (Fig. 6). In the southern tamandua (*T. tetradactyla*), the inactivating mutations of *CHIA5* were identified and the estimated inactivation time of this gene was 6.8 Ma, subsequent to the origin of Vermilingua (34.2 Ma) and after the divergence with the giant anteater (*M. tridactyla*) at 11.3 Ma, suggesting a loss specific to lesser anteaters of the genus *Tamandua* (Emerling et al. 2018). In our study, this gene was not found to be expressed in the salivary glands of the giant anteater. On the other hand, *CHIA5* is functional in insectivorous carnivores (Carnivora) and pangolins (Pholidota), whereas *CHIA1-4* are pseudogenized (Emerling et al. 2018; Tabata et al. 2022). Similar inactivating mutations have been observed in the *CHIA1* gene in carnivores and pangolins and dated to at least 67 Ma, well before the origin of carnivores (46.2 Ma) and pangolins (26.5 Ma) (Emerling et al. 2018). Thus, despite relying on a fully myrmecophagous diet, pangolins have only one functional *CHIA* gene (*CHIA5*), likely due to a historical contingency related to their common inheritance with carnivores (Fig. 6). These analyses have thus revealed contrasting pseudogenization events between convergent myrmecophagous species, with lesser anteaters (genus *Tamandua*) retaining functional orthologs for four out of the five chitin-degrading *CHIA* genes (*CHIA1- 4*), while the Malayan pangolin (*M. javanica*) inherited only the fifth one (*CHIA5*) (Emerling et al. 2018). This peculiar evolutionary history raised the question whether the Malayan pangolin might compensate for the paucity of its functional chitinase gene repertoire by overexpressing *CHIA5* in different digestive organs.

**Figure 6:**
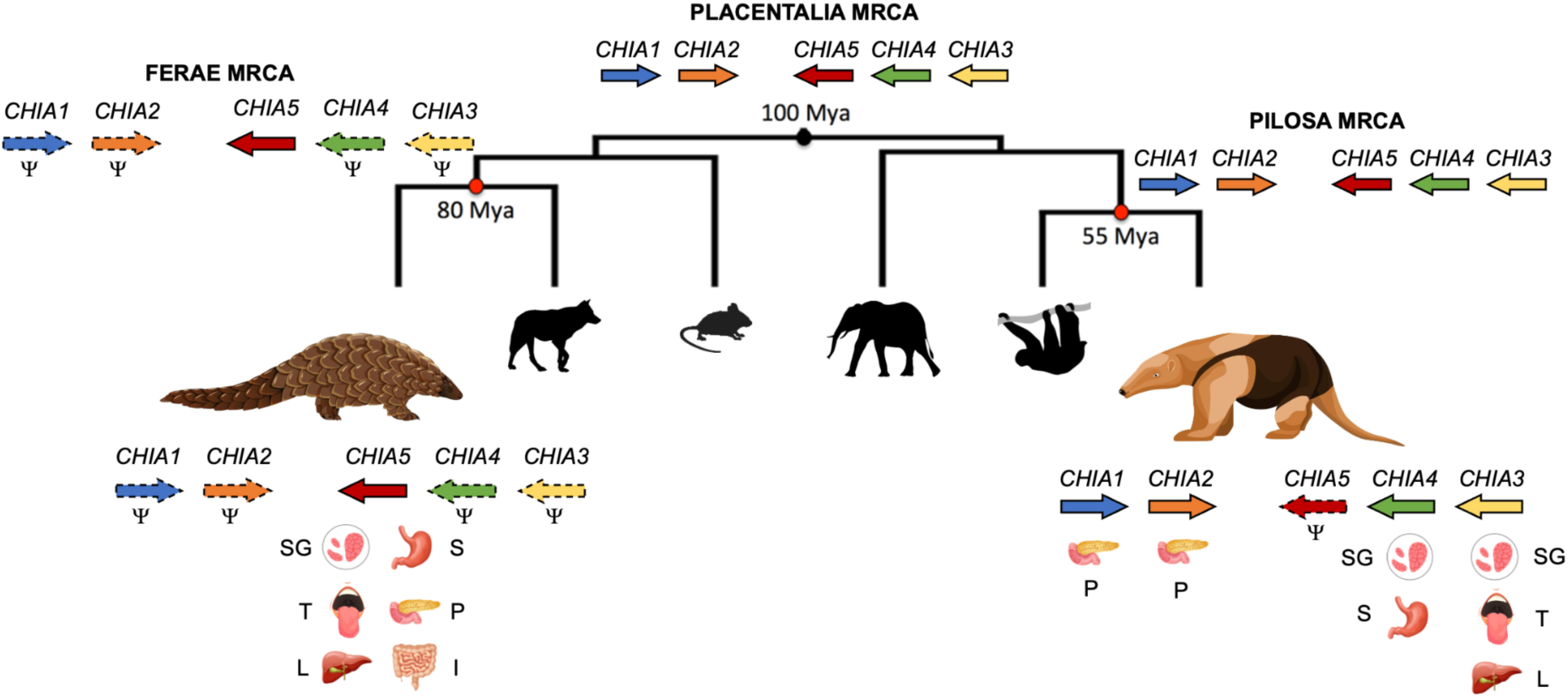
Summary figure presenting the evolution and expression of chitinase acidic (*CHIA*) paralogous genes in the convergently evolved Malayan pangolin (*Manis javanica*) and southern tamandua (*Tamandua tetradactyla*) in their phylogenetic context. Reconstructed *CHIA* gene repertoires are indicated for the two myrmecophagous species and for the most recent common ancestor (MRCA) of placentals, pangolins+carnivores (Ferae) and anteaters+sloths (Pilosa). Non-functional pseudogenes are represented by the Ψ symbol and dashed line contour. Organ icons indicate expression of the corresponding gene in different digestive organs. SG: Salivary glands; S: Stomach; T: Tongue; P: Pancreas; L: Liver; I: Intestine. Silhouettes were obtained from www.phylopic.org and www.vecteezy.com.

Since the presence of enlarged salivary glands is a hallmark of ant-eating mammals, ensuring massive production of saliva to help catch and potentially digest prey, we first investigated chitinase gene expression in mammalian salivary glands. Our comparative transcriptomic study spanning a diversity of species with different diets revealed that, among ant-eating mammals, the Malayan pangolin (*M. javanica*), the southern tamandua (*T. tetradactyla*), and the giant anteater (*M. tridactyla*) all express one or more chitin-degrading genes in their salivary glands. More specifically, we found that *CHIA1* and *CHIA2* were almost never expressed in mammalian salivary glands. By contrast, *CHIA4* was found to be expressed in the giant anteater (*M. tridactyla*) and expression of both *CHIA3* and *CHIA4* was observed in the three southern tamandua (*T. tetradactyla*) individuals surveyed. Moreover, we were able to confirm the hypothesis implying an overexpression of the only functional *CHIA* gene possessed by the Malayan pangolin. Indeed, salivary gland expression profiles of *CHIA5* in *M. javanica* were much higher than in the four other species (Thomas’s nectar bat, mouse, genet, and wild boar) in which we detected expression of this gene, but also substantially higher than the expression of any other chitin-degrading *CHIA* in the 32 other mammalian species considered. Finally, apart from anteaters, *CHIA3* and *CHIA4* were found to be highly expressed only in the two individuals of the insectivorous California leaf-nosed bat (*M. californicus*), but not in any of the other 11 examined bat species, including insectivorous species such as *M. myotis, P. parnellii, and L. evotis*. A possible explanation is that these genes have been pseudogenized in many of these bat species, which would be concordant with the findings of comparative genomic studies reporting widespread pseudogenizations of *CHIA* paralogs across multiple bat species (Emerling et al. 2018), with complete loss of *CHIA1-5* function in non-insectivorous old world fruit bats, most frugivorous bats, and the sanguivorous common vampire bat (Wang et al. 2020). However, although *CHIA4* and *CHIA5* appear to be functional in the insectivorous little brown myotis (*M. lucifugus*; Emerling et al. 2018; Wang et al. 2020), we did not observe expression of these genes in the salivary gland transcriptome we analyzed. Also, *CHIA5* was found to be highly expressed in Thomas’s nectar bat (*H. thomasi*). Although this bat species feeds mostly on nectar and fruits, its diet also includes a substantial part of insects suggesting that CHIA5 might play a role in chitin digestion in the oral cavity, as a result of salivary gland secretion. Transcriptomic analyses of additional digestive tissues besides salivary glands in bats (Vandewege et al. 2020) may further clarify this pattern since chitinolytic activity has previously been reported in the stomachs of seven insectivorous bat species (Strobel et al. 2013). Overall, our chitinase gene expression results therefore support a primary role for salivary glands in prey digestion through the use of distinct *CHIA* paralogs (*CHIA3*, *CHIA4*, and *CHIA5*) in different insect-eating placental mammal species.

Our differential expression comparison of the distinct chitinase paralogs across different organs further highlight the importance of *CHIA5* for Malayan pangolin digestive physiology by confirming its ubiquitous expression in all major tissues of the digestive tract (tongue, salivary glands, stomach, pancreas, liver, and large and small intestines) (Ma et al. 2017; Ma et al. 2019; Cheng et al. 2023; Fig. 6). More specifically, *CHIA5* was found to be expressed at particularly high levels in the stomach and salivary glands. These results are in line with previous proteomic studies that have also identified *CHIA5* as a digestive enzyme (Zhang et al. 2019), which has been confirmed to be highly expressed by RT-qPCR in the specialized oxyntic glands of the stomach (Ma et al. 2018; Cheng et al. 2023), reflecting a key adaptation of the Malayan pangolin to its strictly myrmecophagous diet. By contrast, in the southern tamandua (*T. tetradactyla*) only *CHIA5* is pseudogenized (Emerling et al. 2018; Cheng et al. 2023) and all functional *CHIAs* were found expressed in its digestive tract but not in the same tissues (Fig. 6). *CHIA1* and *CHIA2* were particularly highly expressed in the pancreas whereas *CHIA3* and *CHIA4* were expressed across several other organs of the digestive tract including tongue, salivary glands, stomach, and liver. *CHIA1-4* were also expressed in other non-digestive organs (testes, lungs, and spleen), but their co-expression in the salivary glands of the three southern tamandua individuals sampled here strongly suggests that they play a crucial role in chitin digestion in this myrmecophagous species. Conversely, in the less specialized insectivorous nine-banded armadillo (*D. novemcinctus*), although only *CHIA1* is pseudogenized (Emerling et al. 2018) and therefore not expressed, we did not detect any expression of *CHIA2*, *CHIA3*, and *CHIA4* in the diverse tissues of the individuals studied here, including salivary glands, and *CHIA5* was found only weakly expressed in one spleen sample. Yet, chitinases could still participate in prey digestion in the nine-banded armadillo as they have been isolated from gastric tissues (Smith et al. 1998). We could not confirm this result, given that the liver and colon were the only additional digestive organs besides salivary glands represented in our dataset for this species. However, the comparison with the two myrmecophagous species seems to fit well with its less specialized insectivorous diet and actually further underscores the contrasted specific use of distinct *CHIA* paralogs for chitin digestion in anteaters and pangolins.

Our results demonstrate that in the case of the southern tamandua (*T. tetradactyla*) and the Malayan pangolin (*M. javanica*), two myrmecophagous species that diverged about 100 Ma ago (Meredith et al. 2011), convergent adaptation to myrmecophagy has been achieved in part by using paralogs of different chitinase genes to digest chitin (Fig. 6), probably due to phylogenetic constraints leading to the loss of *CHIA1*, *CHIA2*, *CHIA3*, and *CHIA4* in the most recent common ancestor of Ferae (Carnivora and Pholidota; Emerling et al. 2018). Pangolins and anteaters present extreme morphological adaptations, including the complete loss of dentition, but a detailed study of their feeding apparatus has shown that convergent tooth loss resulted in divergent structures in the internal morphology of their mandible (Ferreira-Cardoso et al. 2019). Our results combined with this observation clearly show that the evolution of convergent phenotypes in myrmecophagous mammals does not necessarily imply similar underlying mechanisms. Our study shows that historical contingency resulted in molecular tinkering (sensu Jacob 1977) of the chitinase gene family at both the genomic and transcriptomic levels in convergently evolved anteaters and pangolins. Working from different starting materials (*i.e.* different *CHIA* paralogs), natural selection led pangolins and anteaters to follow different paths in their convergent adaptation to the myrmecophagous diet.

### Insights from paralogous gene expression in comparative studies

Despite the interest in looking at overall expression patterns to identify the main effect associated with gene expression variation, exploratory comparative transcriptomic analyses have some limitations. Indeed, when comparing the overall gene expression pattern of different species, the first step is to identify comparable elements. These comparable elements can be restricted to single-copy orthologs or extended to homologous gene families containing different paralogs. However, some biases may be introduced during this step (see Li et al. 2023 for a review). On the one hand, focusing only on orthologous genes completely neglects the effects of paralogous gene expression. On the other hand, working with at the scale of large homologous families (orthogroups) often leads to summarizing the expression of multiple orthologous genes into a single expression value. In our case, for example, following the orthogroup detection and summarizing the expression for each orthogroup would have led to a single expression value for the entire chitinase gene family (found as a single orthogroup). By contrast, thanks to our detailed investigation of the evolution of this gene family, phylogenetic and expression analyses of the chitinase orthogroups revealed interesting patterns that would have been missed by the global approach (i.e. effect of contingency bypassed by the relative expression of chitinase family genes). In particular, this approach highlighted differences in gene expression between closely related paralogs (i.e.

*CHIAs*) in the digestive organs of the southern tamandua and the Malayan pangolin, which was crucial for our understanding of the molecular mechanisms involved in this case of convergent dietary adaptation. This result underscores the importance of using both genome- and transcriptome-wide analyses to identify novel candidate genes influencing specific traits, and more targeted approaches based on existing knowledge. The latter is essential to deepen our understanding of the underlying mechanisms observed in specific cases, such as those of convergent evolution linked to historical contingency, as explored in this study.

## Material and Methods

### Chitinase gene family tree reconstruction

#### Reconstruction of chitinase gene family evolution

Mammalian sequences similar to the protein sequence of the human *CHIA* chitinase acidic gene (NP_970615.2) were searched in the NCBI non-redundant protein database using BLASTP (E-value < 10). The protein sequences identified by BLASTP (n = 1,476) were then aligned using MAFFT v7.450 (Katoh and Standley 2013) with the following parameters (--auto --op 1.53 --ep 0.123 --aamatrix BLOSUM62). Preliminary gene trees were then reconstructed with maximum likelihood using RAxML v8.2.11 (Stamatakis 2014) under the LG+G4 model (Le and Gascuel 2008).

From the reconstructed tree, the sequences were filtered according to the following criteria: (1) fast-evolving sequences with a BLAST E-value greater than zero and not belonging to the chitinase family were excluded; (2) in cases of multiple isoforms, only the longest was retained; (3) sequences whose length represented less than at least 50% of the total alignment length were removed; (4) in case of identical sequences of different lengths from the same species the longest was kept; and (5) sequences labeled as “Hypothetical protein” and “Predicted: low quality protein” were discarded. This procedure resulted in a dataset containing 528 mammalian sequences that were realigned using MAFFT with the following parameters (--auto --op 1.53 --ep 0.123 --aamatrix BLOSUM62). This alignment contained 581 amino acid positions and was then cleaned up by removing sites not present in at least 50% of the sequences resulting in a total length of 460 amino acid sites. A maximum likelihood tree was then reconstructed with RAxML-NG v0.9.0 (Kozlov et al. 2019) using 10 tree searches starting from maximum parsimony trees under the LG+G8+F model. The species tree of the 143 mammal species represented in our dataset was reconstructed based on *COI* sequences extracted from the BOLD system database v4 (Ratnasingham and Hebert 2007) by searching for “Chordata” sequences in the “Taxonomy” section. Sequences were aligned using MAFFT with the following parameters (--auto --op 1.53 --ep 0.123 --aamatrix BLOSUM62), the phylogeny was inferred with RAxML under the GTR+G4 model and the topology was then adjusted manually based on the literature to correct ancient relationships. To determine the optimal rooting scheme, a rapid reconciliation between the resulting gene tree and species tree was performed using the TreeRecs reconciliation algorithm based on maximum parsimony (Comte et al. 2020) as implemented in SeaView v5.0.2 (Gouy et al. 2010). The final chitinase gene family tree was produced using the maximum likelihood gene family tree reconciliation approach implemented in GeneRax v.1.1.0 (Morel et al. 2020) using the TreeRecs reconciled tree as input (source and result available from Zenodo).

GeneRax can reconstruct duplications, losses, and horizontal gene transfer events but since the latter are negligible in mammals, only gene duplications and losses have been modeled here (--rec-model UndatedDL) and the LG+G model was used.

#### Ancestral sequence reconstructions

Ancestral sequences of the different paralogs were reconstructed from the reconciled tree using RAxML-NG (--ancestral function --model LG+G8+F). The sequences were then aligned with MAFFT with the following parameters (-- auto --op 1.53 --ep 0.123 --aamatrix BLOSUM62) (source and result files available from Zenodo). Given that active chitinases are characterized by a catalytic site with a conserved amino acid motif (DXXDXDXE; Olland et al. 2009; Hamid et al. 2013), this motif was compared among all available species. Additionally, the six conserved cysteine residues responsible for chitin binding (Tjoelker et al. 2000; Olland et al. 2009) were also investigated.

#### Chitinase gene synteny comparisons

The synteny of the nine chitinase paralogs was compared between the two focal ant-eating species in our global transcriptomic analysis (*T. tetradactyla* and *M. javanica*), an insectivorous xenarthran species (*D. novemcinctus*), an insectivorous primate species with five functional *CHIA* genes (*Carlito syrichta*), and human (*Homo sapiens*). For *H. sapiens*, synteny information was added from Emerling et al. (2018) and completed by using Genomicus v100.01 (Nguyen et al. 2022). For *C. syrichta* and *D. novemcinctus*, genome assemblies were downloaded from the National Center for

Biotechnology Information (NCBI) and from the DNA Zoo (Choo et al. 2016; Dudchenko et al. 2017) for *M. javanica* and *T. tetradactyla*. Synteny information was retrieved by blasting (*megablast*) the different CDS sequences against these assemblies. Scaffold/contig names, positions and direction of BLAST hits were retrieved to compare their synteny (source and result files available from Zenodo). Genes with no significant BLAST hits were considered probably not functional or absent.

### Transcriptome assemblies

#### Salivary gland transcriptomes

Biopsies of submandibular salivary glands (Gil et al. 2018) preserved in RNAlater were obtained from the Mammalian Tissue Collection of the Institut des Sciences de l’Evolution de Montpellier (ISEM) and the JAGUARS collection for 16 individuals representing 12 placental mammal species (Table S1). Total RNA was extracted from individual salivary gland tissue samples using the RNeasy extraction kit (Qiagen, Germany). Then, RNA-seq library construction and Illumina sequencing on a HiSeq 2500 system using paired-end 2x125bp reads were conducted by the Montpellier GenomiX platform (MGX) resulting in 16 newly produced salivary gland transcriptomes. This sampling was completed with the 26 mammalian salivary gland transcriptomes available as paired-end Illumina sequencing reads in the Short Read Archive (SRA) of the NCBI as of December 15th, 2022 representing an additional 21 species (Table S1). This taxon sampling includes representatives from all major mammal superorders Afrotheria (n = 4), Xenarthra (n = 4), Euarchontoglires (n = 4), and Laurasiatheria (n = 21) and covers six different diet categories: carnivory (n = 4), frugivory and herbivory (n = 8), insectivory (n = 9), myrmecophagy (n = 5), and omnivory (n = 7) (Table S1). Four of the five lineages in which myrmecophagous mammals evolved are represented: southern aardwolf (*P. cristatus*, Carnivora), Malayan pangolin (*M. javanica*, Pholidota), southern naked-tailed armadillo (*C. unicinctus*, Cingulata), giant anteater (*M. tridactyla*, Pilosa), and southern tamandua (*T. tetradactyla*, Pilosa). Species replicates in the form of different individuals were included for the southern tamandua (*T. tetradactyla*; n = 3), the nine-banded armadillo (*D. novemcinctus*; n = 3), the Malayan pangolin (*M. javanica*; n = 2), the vampire bat (*Desmodus rotundus*; n = 2), and the California leaf-nosed bat (*Macrotus californicus*; n = 2). We unfortunately were not able to obtain fresh salivary gland samples from the aardvark (*O. afer*, Tubulidentata), the only missing myrmecophagous lineage in our sampling.

#### Transcriptomes from additional organs

Tissue biopsies from nine additional organs (testis, lungs, heart, spleen, tongue, pancreas, stomach, liver, and small intestine) were sampled during dissections of three roadkill individuals of southern tamandua (*T. tetradactyla*; Table S1). Total RNA extractions from these RNAlater-preserved tissues, RNA-seq library construction, and sequencing were conducted as described above resulting in 13 newly generated transcriptomes. For comparative purposes, 21 additional transcriptomes of nine-banded armadillo (*D. novemcinctus*) representing eight organs and 32 transcriptomes of Malayan pangolin (*M. javanica*) representing 16 organs were downloaded from SRA (Table S1).

### Comparative transcriptomics

*Transcriptome assemblies and quality control* - Adapters and low quality reads were removed from raw sequencing data using fastp v0.19.6 (Chen et al. 2018). Reads were allowed a minimum of 40% of bases with a PHRED score at least 15 (-- qualified_quality_phred ≥ 15), as suggested by (MacManes 2014). Then, *de novo* assembly was performed on each individual transcriptome sample using Trinity v2.8.4 (Grabherr et al. 2011) using cleaned paired-end reads (--seqType fq --left R1.fastq --right R2.fastq; result files available from Zenodo). For one individual vampire bat (*D. rotundus*), three salivary gland transcriptomes (SRR606902, SRR606908, and SRR606911) were combined to obtain a better assembly. For each of the 104 transcriptome assemblies, completeness was assessed by the presence of Benchmark Universal Single Copy Orthologs (BUSCO v5) based on a predefined dataset (mammalia_odb10) of 9,226 single-copy orthologs conserved in over 90% of mammalian species (Manni et al. 2021). This pipeline was run through the gVolante web server (Nishimura et al. 2017) to evaluate the percentage of complete, duplicated, fragmented and missing single copy orthologs within each transcriptome (Table S2).

#### Transcriptome annotation and orthogroup inference

The 104 transcriptome assemblies were annotated following the pipeline implemented in assembly2ORF (https://github.com/ellefeg/assembly2orf). This pipeline combines evidence-based and gene-model-based predictions. First, potential transcripts of protein-coding genes are extracted based on similarity searches (BLAST) against the peptides of Metazoa found in Ensembl (Yates et al. 2020). Then, using both protein similarity and exonerate functions (Slater and Birney 2005), a frameshift correction is applied to candidate transcripts. Candidate open reading frames (ORFs) are predicted using TransDecoder (https://github.com/TransDecoder/TransDecoder) and annotated based on homology information inferred from both BLAST and Hmmscan searches. Finally, to be able to compare the transcriptomes obtained from all species, we relied on the inference of gene orthogroups. The orthogroup inference for the translated candidate ORFs was performed using OrthoFinder v2 (Emms and Kelly 2019) using FastTree (Price et al. 2010) for gene tree reconstructions. For expression analyses, orthogroups containing more than 20 copies for at least one species were discarded, resulting in the selection of 13,392 orthogroups for further analyses.

#### Gene expression analyzes

Quantification of transcript expression was performed on Trinity assemblies with Kallisto v.0.46.1 (Bray et al. 2016) using the *align_and_estimate_abundance.pl* script provided in the Trinity suite (Grabherr et al. 2011). Kallisto relies on pseudo-alignments of the reads to search for the original transcript of a read without looking for a perfect alignment (as opposed to classical quantification by counting the reads aligned on the assembled transcriptome; Wolf 2013). Counts (raw number of mapped reads) and the Transcripts Per kilobase Million are reported (result files available from Zenodo). Based on the previously inferred orthogroups, orthogroup-level abundance estimates were imported and summarized using tximport (Soneson et al. 2016). To minimize sequencing depth variation across samples and gene outlier effect (a few highly and differentially expressed genes may have strong and global influence on every gene read count), orthogroup-level raw reads counts were normalized using the median of the ratios of observed counts using DESeq2 (Love et al. 2014).

#### Chitinase expression in salivary glands

The chitinase orthogroup was extracted from the orthogroups inferred by OrthoFinder2 using BLASTX with the reference chitinase database previously created. The 476 amino acid sequences composing this orthogroup were assigned to the nine chitinase orthologs (*CHIA1-5*, *CHIT1*, *CHI3L1*, *CHI3L2*, *OVGP1*) using the maximum likelihood Evolutionary Placement Algorithm implemented in RAxML-EPA (Berger et al. 2011) with the reference chitinase sequence alignment and reconciled phylogenetic tree previously inferred using GeneRax (result files available from Zenodo).

This allowed excluding three additional contaminant sequences and dividing the chitinase orthogroup into nine sub-orthogroups corresponding to each chitinase paralog. To take advantage of the transcriptome-wide expression information for the expression standardization, these new orthogroups were included in the previous orthogroup-level abundance matrix estimates and the same normalization approach using DESeq2 was conducted. Finally, gene-level abundance estimates for all chitinase paralogs were extracted and compared on a log10 scale.

## Data and Resource Availability

Raw RNAseq Illumina reads have been submitted to the Short Read Archive (SRA) of the National Center for Biotechnology Information (NCBI) and are available under BioProject number PRJNA909065. Transcriptome assemblies, phylogenetic datasets, corresponding trees, and other supplementary materials are available from zenodo.org (https://doi.org/10.5281/zenodo.7355329).

## Acknowledgments

We would like to thank Hugues Parrinello (Montpellier GenomiX platform) for advice on RNAseq, Mariana Escobar Rodríguez and Gautier Debaecker for help with transcriptome assembly and annotation, and Marie Sémon for providing useful advice on RNAseq statistical analyses. We are also indebted to Frank Knight, Mark Scherz, Miguel Vences, Andolalao Rakotoarison, Nico Avenant, Pierre-Henri Fabre, Quentin Martinez, Nathalie Delsuc, Aude Caizergues, Roxanne Schaub, Lionel Hautier, Fabien Condamine, Sérgio Ferreira-Cardoso, and François Catzeflis for their help with tissue sampling. We also thank the three anonymous referees for their helpful comments. Computational analyses benefited from the Montpellier Bioinformatics Biodiversity (MBB) platform. The JAGUARS collection is supported through a grants from the Collectivité Territoriale de Guyane, from the European Union, and from Direction Générale des Territoires et de la Mer / Préfet de la Région Guyane attributed to Kwata NGO. This work has been supported by grants from the European Research Council (ConvergeAnt project: ERC-2015-CoG-683257) and Investissements d’Avenir of the Agence Nationale de la Recherche (CEBA: ANR-10-LABX-25-01; CEMEB: ANR-10-LABX-0004). This is contribution ISEM 2024-XXX of the Institut des Sciences de l’Evolution de Montpellier.

## Notes

https://doi.org/10.5281/zenodo.7355329

### Competing Interest Statement

The authors have declared no competing interest.

### Summary of Updates

This version of the manuscript has been revised according to reviewers' comments. The title has been changed, the introduction completely rewritten, and two new figures have been added.

