## Supplementary material for "Transcriptomic data reveal divergent paths of chitinase evolution underlying dietary convergence in anteaters and pangolins": Table S1

| Sample name | Species | Tissue type | Individual name | Sex | Source | Country of origin | Study |
|---|---|---|---|---|---|---|---|
| CABuniCU04 | *Cabassous unicinctus* | Salivary gland (submandibular) | M2757 | Male | JAGUARS | French Guiana | This study |
| SRR5889344 | *Canis lupus familiaris* | Salivary gland | NA | NA | SRA NCBI | USA | Broad Institute, unpublished |
| SRR7064957 | *Carollia sowelli* | Salivary gland (submandibular) | TX169114 | Male | SRA NCBI | Guatemala | Vandewege *et al* , 2020 |
| SRR7064954 | *Centurio senex* | Salivary gland (submandibular) | TK169257 | Female | SRA NCBI | Guatemala | Vandewege *et al* , 2020 |
| DASnovFK06C | *Dasypus novemcinctus* | Salivary gland (submandibular) | FK06 | Female | ISEM | USA | This study |
| SRR494777 | *Dasypus novemcinctus* | Cerebellum | 0986 | Male | SRA NCBI | USA | Broad Institute, unpublished |
| SRR494780 | *Dasypus novemcinctus* | Cerebellum | 0986 | Male | SRA NCBI | USA | Broad Institute, unpublished |
| SRR494772 | *Dasypus novemcinctus* | Colon | 0986 | Male | SRA NCBI | USA | Broad Institute, unpublished |
| SRR494774 | *Dasypus novemcinctus* | Colon | 0986 | Male | SRA NCBI | USA | Broad Institute, unpublished |
| SRR494769 | *Dasypus novemcinctus* | Heart | 0986 | Male | SRA NCBI | USA | Broad Institute, unpublished |
| SRR494773 | *Dasypus novemcinctus* | Heart | 0986 | Male | SRA NCBI | USA | Broad Institute, unpublished |
| SRR6206903 | *Dasypus novemcinctus* | Heart | NA | NA | SRA NCBI | USA | Chen *et al*, 2019 |
| SRR494775 | *Dasypus novemcinctus* | Kidney | 0986 | Male | SRA NCBI | USA | Broad Institute, unpublished |
| SRR494779 | *Dasypus novemcinctus* | Kidney | 0986 | Male | SRA NCBI | USA | Broad Institute, unpublished |
| SRR6206908 | *Dasypus novemcinctus* | Kidney | NA | NA | SRA NCBI | USA | Chen *et al*, 2019 |
| SRR494766 | *Dasypus novemcinctus* | Liver | 0986 | Male | SRA NCBI | USA | Broad Institute, unpublished |
| SRR494778 | *Dasypus novemcinctus* | Liver | 0986 | Male | SRA NCBI | USA | Broad Institute, unpublished |
| SRR6206913 | *Dasypus novemcinctus* | Liver | NA | NA | SRA NCBI | USA | Chen *et al*, 2019 |
| SRR494776 | *Dasypus novemcinctus* | Lung | 0986 | Male | SRA NCBI | USA | Broad Institute, unpublished |
| SRR494781 | *Dasypus novemcinctus* | Lung | 0986 | Male | SRA NCBI | USA | Broad Institute, unpublished |
| SRR6206918 | *Dasypus novemcinctus* | Lung | NA | NA | SRA NCBI | USA | Chen *et al*, 2019 |
| SRR494770 | *Dasypus novemcinctus* | Muscle | 0986 | Male | SRA NCBI | USA | Broad Institute, unpublished |
| SRR494771 | *Dasypus novemcinctus* | Muscle | 0986 | Male | SRA NCBI | USA | Broad Institute, unpublished |
| SRR6206923 | *Dasypus novemcinctus* | Muscle | NA | NA | SRA NCBI | USA | Chen *et al*, 2019 |
| DASnovFK06A | *Dasypus novemcinctus* | Salivary gland (submandibular) | FK06 | Female | ISEM | USA | This study |
| DASnovFK08A | *Dasypus novemcinctus* | Salivary gland (submandibular) | FK08 | Male | ISEM | USA | This study |
| SRR494767 | *Dasypus novemcinctus* | Spleen | 0986 | Male | SRA NCBI | USA | Broad Institute, unpublished |
| SRR494768 | *Dasypus novemcinctus* | Spleen | 0986 | Male | SRA NCBI | USA | Broad Institute, unpublished |
| SRR606902 | *Desmodus rotundus* | Salivary gland | NA | NA | SRA NCBI | Brazil | National Institute of Allergy and Infectious Diseases, unpublished |
| SRR606908 | *Desmodus rotundus* | Salivary gland | NA | NA | SRA NCBI | Brazil | National Institute of Allergy and Infectious Diseases, unpublished |
| SRR606911 | *Desmodus rotundus* | Salivary gland | NA | NA | SRA NCBI | Brazil | National Institute of Allergy and Infectious Diseases, unpublished |
| SRR7064949 | *Desmodus rotundus* | Salivary gland (submandibular) | TK169403 | Male | SRA NCBI | Guatemala | Vandewege *et al* , 2020 |
| ELEmyuNA02 | *Elephantulus myurus* | Salivary gland (submandibular) | TDR | NA | ISEM | South Africa | This study |
| SRR19501516 | *Elephas maximus* | Salivary gland | 445 | Female | SRA NCBI | Germany | Leibniz Institute for Zoo and Wildlife Research, unpublished |
| ERIeurRA02 | *Erinaceus europaeus* | Salivary gland (submandibular) | RA03 | NA | ISEM | France | This study |
| SRR3218717 | *Felis catus* | Salivary gland | NA | Female | SRA NCBI | USA | Visser *et al* , 2019 |
| GENgenRA01 | *Genetta genetta* | Salivary gland (submandibular) | RA02 | NA | ISEM | France | This study |
| SRR7064959 | *Glossophaga commissarisi* | Salivary gland (submandibular) | TK169022 | Male | SRA NCBI | Guatemala | Vandewege *et al* , 2020 |
| SRR1957200 | *Homo sapiens* | Salivary gland | NA | NA | SRA NCBI | USA | Duff *et al*, 2015 |
| SRR7064953 | *Hsunycteris tomasi* | Salivary gland (submandibular) | TK104601 | Male | SRA NCBI | Equator | Vandewege *et al* , 2020 |
| SRR7064956 | *Lophostoma evotis* | Salivary gland (submandibular) | TK169173 | Female | SRA NCBI | Guatemala | Vandewege *et al* , 2020 |
| SRR1023040 | *Macrotus californicus* | Salivary gland (submandibular) | NA | Male | SRA NCBI | USA | Texas Tech University; unpublished |
| SRR7064950 | *Macrotus californicus* | Salivary gland (submandibular) | TK163824 | Male | SRA NCBI | USA | Vandewege *et al* , 2020 |
| SRR2561209 | *Manis javanica* | Brain | NA | Female | SRA NCBI | Malaysia | Yusoff *et al*, 2016 |
| SRR2547558 | *Manis javanica* | Cerebellum | NA | Female | SRA NCBI | Malaysia | Yusoff *et al*, 2016 |
| SRR2561211 | *Manis javanica* | Heart | NA | Female | SRA NCBI | Malaysia | Yusoff *et al*, 2016 |
| SRR2561212 | *Manis javanica* | Kidney | NA | Female | SRA NCBI | Malaysia | Yusoff *et al*, 2016 |
| SRR7641089 | *Manis javanica* | Large intestine | NA | Female | SRA NCBI | China | Ma *et al*, 2019 |
| SRR7641090 | *Manis javanica* | Large intestine | NA | Female | SRA NCBI | China | Ma *et al*, 2019 |
| SRR8943550 | *Manis javanica* | Large intestine | MJ63 | Female | SRA NCBI | China | Hu *et al* , 2020 |
| SRR2561213 | *Manis javanica* | Liver | NA | Female | SRA NCBI | Malaysia | Yusoff *et al*, 2016 |
| SRR5341161 | *Manis javanica* | Liver | NA | Female | SRA NCBI | China | Ma *et al*, 2017 |
| SRR7641080 | *Manis javanica* | Liver | NA | Female | SRA NCBI | China | Ma *et al*, 2019 |
| SRR7641081 | *Manis javanica* | Liver | NA | Female | SRA NCBI | China | Ma *et al*, 2019 |
| SRR7641087 | *Manis javanica* | Liver | NA | Female | SRA NCBI | China | Ma *et al*, 2019 |
| SRR7641088 | *Manis javanica* | Liver | NA | Female | SRA NCBI | China | Ma *et al*, 2019 |
| SRR2561214 | *Manis javanica* | Lung | NA | Female | SRA NCBI | Malaysia | Yusoff *et al*, 2016 |
| SRR8943548 | *Manis javanica* | Lung | MJ63 | Female | SRA NCBI | China | Hu *et al* , 2020 |
| SRR5837767 | *Manis javanica* | Muscle | NA | NA | SRA NCBI | China | Jiangsu Normal University; unpublished |
| SRR8943546 | *Manis javanica* | Ovary | MJ63 | Female | SRA NCBI | China | Hu *et al* , 2020 |
| SRR7641079 | *Manis javanica* | Pancreas | NA | Female | SRA NCBI | China | Ma *et al*, 2019 |
| SRR7641082 | *Manis javanica* | Pancreas | NA | Female | SRA NCBI | China | Ma *et al*, 2019 |
| SRR8943545 | *Manis javanica* | Pancreas | MJ63 | Female | SRA NCBI | China | Hu *et al* , 2020 |
| SRR5337837 | *Manis javanica* | Salivary gland | NA | Female | SRA NCBI | China | Ma *et al*, 2017 |
| SRR7641084 | *Manis javanica* | Salivary gland | NA | Female | SRA NCBI | China | Ma *et al*, 2019 |
| SRR3923846 | *Manis javanica* | Skin | NA | Female | SRA NCBI | Malaysia | Yusoff *et al*, 2016 |
| SRR5328124 | *Manis javanica* | Small intestine | NA | Female | SRA NCBI | China | Ma *et al*, 2017 |
| SRR8943543 | *Manis javanica* | Small intestine | MJ63 | Female | SRA NCBI | China | Hu *et al* , 2020 |
| SRR2561215 | *Manis javanica* | Spleen | NA | Female | SRA NCBI | Malaysia | Yusoff *et al*, 2016 |
| SRR7641085 | *Manis javanica* | Stomach | NA | Female | SRA NCBI | China | Ma *et al*, 2019 |
| SRR7641086 | *Manis javanica* | Stomach | NA | Female | SRA NCBI | China | Ma *et al*, 2019 |
| SRR8943542 | *Manis javanica* | Stomach | MJ63 | Female | SRA NCBI | China | Hu *et al* , 2020 |
| SRR2561216 | *Manis javanica* | Thymus | NA | Female | SRA NCBI | Malaysia | Yusoff *et al*, 2016 |
| SRR7641083 | *Manis javanica* | Tongue | NA | Female | SRA NCBI | China | Ma *et al*, 2019 |
| SRR8943544 | *Manis javanica* | Tongue | MJ63 | Female | SRA NCBI | China | Hu *et al* , 2020 |
| MELmelRA01 | *Meles meles* | Salivary gland (submandibular) | RA01 | Female | ISEM | France | This study |
| MICspMV01 | *Microgale brevicaudata* | Salivary gland (submandibular) | MV03 | NA | ISEM | Madagascar | This study |
| SRR5878900 | *Mus musculus* | Salivary gland | NA | NA | SRA NCBI | USA | Metwalli *et al*, 2018 |
| MYOcoyPH03 | *Myocastor coypus* | Salivary gland (submandibular) | Myo2 | NA | ISEM | France | This study |
| SRR7064951 | *Myotis lucifugus* | Salivary gland (submandibular) | TK125917 | Male | SRA NCBI | USA | Vandewege *et al* , 2020 |
| MYRtriCAY01 | *Myrmecophaga tridactyla* | Salivary gland (submandibular) | M3023 | Male | JAGUARS | French Guiana | This study |
| ERR2076303 | *Ovis aries* | Salivary gland | NA | Female | SRA NCBI | USA | Clark *et al*, 2017 |
| PROcri01 | *Proteles cristatus* | Salivary gland (submandibular) | TS307 | Male | ISEM | South Africa | This study |
| SRR7064955 | *Pteronotus parnellii* | Salivary gland (submandibular) | TK169174 | Female | SRA NCBI | Guatemala | Vandewege *et al* , 2020 |
| SRR3056926 | *Rattus norvegicus* | Salivary gland | NA | NA | SRA NCBI | USA | Barasch *et al*, 2017 |
| SRR7064958 | *Sturnira hondurensis* | Salivary gland (submandibular) | TK169029 | Male | SRA NCBI | Guatemala | Vandewege *et al* , 2020 |
| SRR5802558 | *Sus scrofa* | Salivary gland | NA | Male | SRA NCBI | China | China Agricultural Univeristy ; unpublished |
| TAMtetTT78 | *Tamandua tetradactyla* | Glandular stomach | M3075 | Male | JAGUARS | French Guiana | This study |
| TAMtetTT75 | *Tamandua tetradactyla* | Heart | M3075 | Male | JAGUARS | French Guiana | This study |
| TAMtetTT59 | *Tamandua tetradactyla* | Liver | M3075 | Male | JAGUARS | French Guiana | This study |
| TAMtetTT73 | *Tamandua tetradactyla* | Lung | M3075 | Male | JAGUARS | French Guiana | This study |
| TAMtetTT79 | *Tamandua tetradactyla* | Muscular stomach | M3075 | Male | JAGUARS | French Guiana | This study |
| TAMtetTT67 | *Tamandua tetradactyla* | Pancreas | M3075 | Male | JAGUARS | French Guiana | This study |
| TAMtetB01 | *Tamandua tetradactyla* | Salivary gland (submandibular) | M2813 | Male | JAGUARS | French Guiana | This study |
| TAMtetFC01 | *Tamandua tetradactyla* | Salivary gland (submandibular) | T7380 | Female | ISEM | French Guiana | This study |
| TAMtetTT49 | *Tamandua tetradactyla* | Salivary gland (submandibular) | M3075 | Male | JAGUARS | French Guiana | This study |
| TAMtetTT99 | *Tamandua tetradactyla* | Small intestine | M3075 | Male | JAGUARS | French Guiana | This study |
| TAMterTT62 | *Tamandua tetradactyla* | Spleen | M3075 | Male | JAGUARS | French Guiana | This study |
| TAMtetFC04 | *Tamandua tetradactyla* | Spleen | T7380 | Female | ISEM | French Guiana | This study |
| TAMtetR05 | *Tamandua tetradactyla* | Testis | M2813 | Male | JAGUARS | French Guiana | This study |
| TAMtetTT70 | *Tamandua tetradactyla* | Testis | M3075 | Male | JAGUARS | French Guiana | This study |
| TAMtetB07 | *Tamandua tetradactyla* | Tongue | M2813 | Male | JAGUARS | French Guiana | This study |
| TAMtetTT55 | *Tamandua tetradactyla* | Tongue | M3075 | Male | JAGUARS | French Guiana | This study |
| SETsetMV01 | *Tenrec ecaudatus* | Salivary gland (submandibular) | MV01 | NA | ISEM | Madagascar | This study |
| SRR7064952 | *Trachops cirrhosus* | Salivary gland (submandibular) | TK167830 | Male | SRA NCBI | Costa Rica | Vandewege *et al* , 2020 |
| SRR1663490 | *Uroderma bilobatum* | Salivary gland (submandibular) | NA | Male | SRA NCBI | Uruguay | Feijoo *et al*, 2017 |
