## Supplementary material for "Transcriptomic data reveal divergent paths of chitinase evolution underlying dietary convergence in anteaters and pangolins": Table S2

| Sample | Complete BUSCOs (C) | Fragmented BUSCOs (F) | Missing BUSCOs (M) |
|---|---|---|---|
| Tamandua_tetradactyla_M3075_Pancreas | 1643 | 730 | 6853 |
| Pteronotus_parnellii_SRR7064955_SalivaryGland | 1831 | 933 | 6462 |
| Dasypus_novemcinctus_SRR494773_Heart | 2478 | 790 | 5958 |
| Dasypus_novemcinctus_SRR494769_Heart | 2507 | 726 | 5993 |
| Proteles_cristatus_SalivaryGland_TS307 | 2856 | 783 | 5587 |
| Sturnira_hondurensis_SRR7064958_SalivaryGland | 2872 | 881 | 5473 |
| Macrotus_californicus_SRR1023040_SalivaryGland | 2953 | 823 | 5450 |
| Centurio_senex_SRR7064954_SalivaryGland | 3001 | 899 | 5326 |
| Uroderma_bilobatum_SRR1663490_SalivaryGland | 3134 | 702 | 5390 |
| Dasypus_novemcinctus_SRR494779_Kidney | 3144 | 778 | 5304 |
| Dasypus_novemcinctus_SRR494775_Kidney | 3195 | 772 | 5259 |
| Glossophaga_commissarisi_SRR7064959_SalivaryGland | 3210 | 852 | 5164 |
| Tamandua_tetradactyla_T7380_Spleen | 3303 | 912 | 5011 |
| Dasypus_novemcinctus_SRR494770_Muscle | 3305 | 713 | 5208 |
| Hsunycteris_tomasi_SRR7064953_SalivaryGland | 3337 | 1030 | 4859 |
| Dasypus_novemcinctus_SRR494771_Muscle | 3374 | 714 | 5138 |
| Carollia_sowelli_SRR7064957_SalivaryGland | 3390 | 821 | 5015 |
| Erinaceus_europaeus_RA03_SalivaryGland | 3432 | 782 | 5012 |
| Trachops_cirrhosus_SRR7064952_SalivaryGland | 3507 | 733 | 4986 |
| Dasypus_novemcinctus_SRR6206903_Heart | 3595 | 739 | 4892 |
| Lophostoma_evotis_SRR7064956_SalivaryGland | 3616 | 839 | 4771 |
| Tamandua_tetradactyla_T7380_SalivaryGland | 3650 | 697 | 4879 |
| Dasypus_novemcinctus_SRR494766_Liver | 3714 | 755 | 4757 |
| Dasypus_novemcinctus_SRR494778_Liver | 3741 | 732 | 4753 |
| Desmodus_rotundus_SRR7064949_SalivaryGland | 4043 | 673 | 4510 |
| Myocastor_coypus_Myo2_SalivaryGland | 4043 | 694 | 4489 |
| Tamandua_tetradactyla_M3075_Heart | 4061 | 898 | 4267 |
| Sus_scrofa_SRR5802558_SalivaryGland | 4188 | 868 | 4170 |
| Dasypus_novemcinctus_SRR6206908_Kidney | 4220 | 755 | 4251 |
| Ovis_aries_ERR2076303_SalivaryGland | 4335 | 778 | 4113 |
| Manis_javanica_SRR5837767_Muscle | 4335 | 727 | 4164 |
| Dasypus_novemcinctus_SRR6206923_Muscle | 4397 | 668 | 4161 |
| Myrmecophaga_tridactyla_M3023_SalivaryGland | 4424 | 892 | 3910 |
| Microgale_brevicaudata_MV03_SalivaryGland | 4477 | 713 | 4036 |
| Dasypus_novemcinctus_FK06C_SalivaryGland | 4578 | 637 | 4011 |
| Macrotus_californicus_SRR7064950_SalivaryGland | 4581 | 740 | 3905 |
| Manis_javanica_SRR7641086_Stomach | 4582 | 827 | 3817 |
| Cabassous_unicinctus_M2757_SalivaryGland | 4647 | 976 | 3603 |
| Manis_javanica_SRR5328124_SmallIntestine | 4725 | 719 | 3782 |
| Dasypus_novemcinctus_SRR6206913_Liver | 4776 | 623 | 3827 |
| Homo_sapiens_SRR1957200_SalivaryGland | 4779 | 961 | 3486 |
| Myotis_lucifugus_SRR7064951_SalivaryGland | 4813 | 861 | 3552 |
| Tamandua_tetradactyla_M3075_Liver | 4824 | 782 | 3620 |
| Manis_javanica_SRR7641085_Stomach | 4843 | 760 | 3623 |
| Dasypus_novemcinctus_SRR494774_Colon | 4848 | 709 | 3669 |
| Dasypus_novemcinctus_SRR494777_Cerebellum | 4905 | 778 | 3543 |
| Dasypus_novemcinctus_SRR494772_Colon | 4906 | 726 | 3594 |
| Dasypus_novemcinctus_FK06A_SalivaryGland | 4908 | 621 | 3697 |
| Genetta_genetta_RA02_SalivaryGland | 4926 | 685 | 3615 |
| Dasypus_novemcinctus_SRR494780_Cerebellum | 4964 | 753 | 3509 |
| Dasypus_novemcinctus_FK08_SalivaryGland | 5021 | 606 | 3599 |
| Manis_javanica_SRR7641084_SalivaryGland | 5063 | 742 | 3421 |
| Elephantulus_myurus_TDR_SalivaryGland | 5122 | 570 | 3534 |
| Tamandua_tetradactyla_M2813_Tongue | 5137 | 744 | 3345 |
| Manis_javanica_SRR5341161_Liver | 5160 | 575 | 3491 |
| Dasypus_novemcinctus_SRR494781_Lungs | 5164 | 775 | 3287 |
| Tamandua_tetradactyla_M2813_SalivaryGland | 5172 | 482 | 3572 |
| Tamandua_tetradactyla_M3075_SmallIntestine | 5192 | 942 | 3092 |
| Dasypus_novemcinctus_SRR494776_Lungs | 5205 | 741 | 3280 |
| Tamandua_tetradactyla_M3075_GlandularStomach | 5224 | 977 | 3025 |
| Manis_javanica_SRR7641081_Liver | 5287 | 662 | 3277 |
| Tamandua_tetradactyla_M3075_MuscularStomach | 5329 | 725 | 3172 |
| Tamandua_tetradactyla_M3075_Tongue | 5334 | 927 | 2965 |
| Manis_javanica_SRR7641087_Liver | 5361 | 627 | 3238 |
| Manis_javanica_SRR7641089_LargeIntestine | 5382 | 762 | 3082 |
| Manis_javanica_SRR5337837_SalivaryGland | 5397 | 661 | 3168 |
| Meles_meles_RA01_SalivaryGland | 5412 | 507 | 3307 |
| Manis_javanica_SRR7641090_LargeIntestine | 5415 | 723 | 3088 |
| Desmodus_rotundus_SRR606902_908_911_SalivaryGland | 5469 | 496 | 3261 |
| Tamandua_tetradactyla_M3075_Spleen | 5501 | 839 | 2886 |
| Felis_catus_SRR3218717_SalivaryGland | 5587 | 723 | 2916 |
| Dasypus_novemcinctus_SRR494768_Spleen | 5601 | 621 | 3004 |
| Dasypus_novemcinctus_SRR494767_Spleen | 5614 | 630 | 2982 |
| Manis_javanica_SRR7641088_Liver | 5625 | 499 | 3102 |
| Manis_javanica_SRR3923846_Skin | 5651 | 613 | 2962 |
| Manis_javanica_SRR7641080_Liver | 5662 | 502 | 3062 |
| Manis_javanica_SRR2561213_Liver | 5715 | 625 | 2886 |
| Manis_javanica_SRR7641083_Tongue | 5739 | 581 | 2906 |
| Tamandua_tetradactyla_M3075_SalivaryGland | 5767 | 490 | 2969 |
| Manis_javanica_SRR2561211_Heart | 5788 | 545 | 2893 |
| Canis_lupus_familiaris_SRR5889344_SalivaryGland | 5820 | 494 | 2912 |
| Manis_javanica_SRR7641082_Pancreas | 5872 | 622 | 2732 |
| Elephas_maximus_SRR19501516_SalivaryGland | 5921 | 623 | 2682 |
| Manis_javanica_SRR8943546_Ovary | 5958 | 528 | 2740 |
| Rattus_norvegicus_SRR3056926_SalivaryGland | 6030 | 213 | 2983 |
| Tenrec_ecaudatus_MV01_SalivaryGland | 6049 | 565 | 2612 |
| Manis_javanica_SRR8943544_Tongue | 6075 | 573 | 2578 |
| Manis_javanica_SRR2561212_Kidney | 6116 | 635 | 2475 |
| Dasypus_novemcinctus_SRR6206918_Lungs | 6123 | 583 | 2520 |
| Manis_javanica_SRR8943542_Stomach | 6128 | 442 | 2656 |
| Tamandua_tetradactyla_M2813_Testis | 6161 | 692 | 2373 |
| Manis_javanica_SRR7641079_Pancreas | 6203 | 582 | 2441 |
| Tamandua_tetradactyla_M3075_Lungs | 6242 | 813 | 2171 |
| Manis_javanica_SRR8943550_LargeIntestine | 6293 | 457 | 2476 |
| Mus_musculus_SRR5878900_SalivaryGland | 6414 | 448 | 2364 |
| Manis_javanica_SRR8943545_Pancreas | 6458 | 440 | 2328 |
| Manis_javanica_SRR2547558_Cerebellum | 6528 | 644 | 2054 |
| Manis_javanica_SRR2561215_Spleen | 6541 | 578 | 2107 |
| Manis_javanica_SRR8943548_Lung | 6547 | 475 | 2204 |
| Manis_javanica_SRR8943543_SmallIntestine | 6550 | 536 | 2140 |
| Manis_javanica_SRR2561216_Thymus | 6657 | 628 | 1941 |
| Manis_javanica_SRR2561214_Lungs | 6803 | 571 | 1852 |
| Manis_javanica_SRR2561209_Cerebrum | 6823 | 611 | 1792 |
| Tamandua_tetradactyla_M3075_Testis | 7068 | 613 | 1545 |

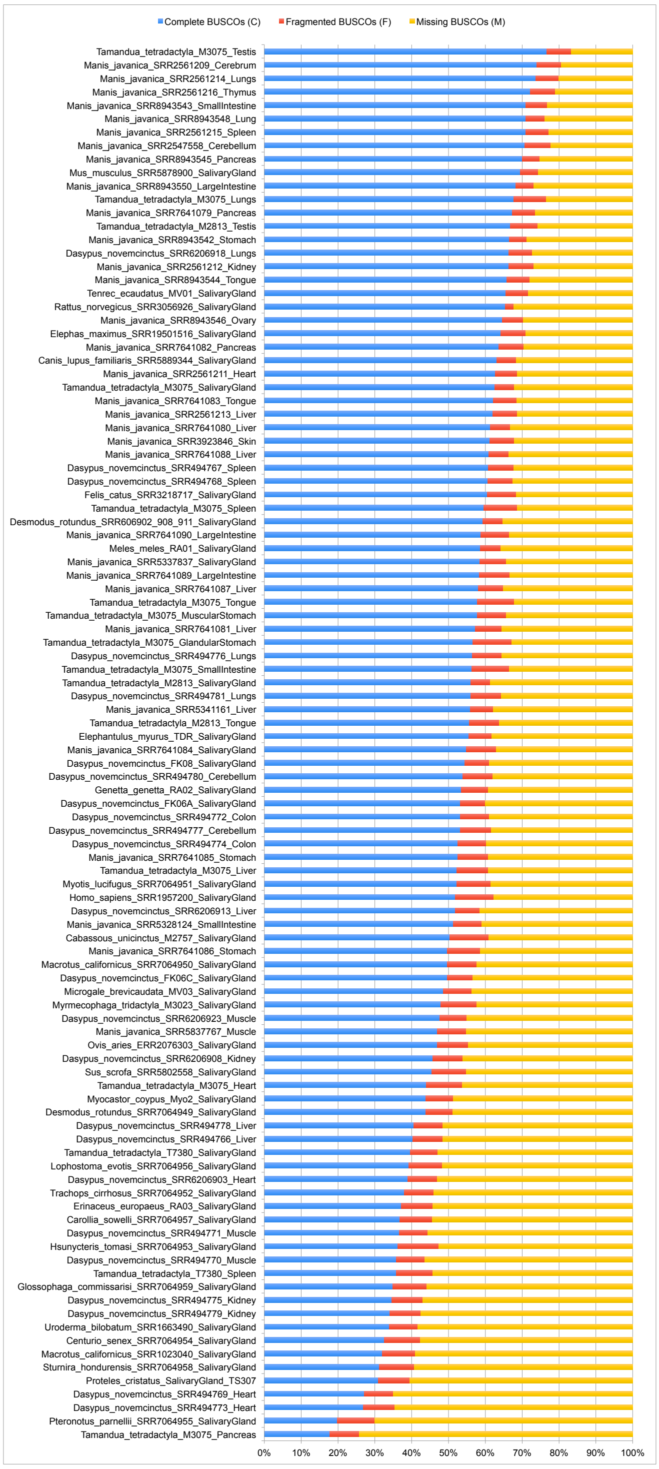
